# Hepatic mTORC2 compensates for loss of adipose mTORC2 in mediating energy storage and glucose homeostasis

**DOI:** 10.1101/2022.12.21.521172

**Authors:** Irina C. Frei, Diana Weissenberger, Christoph Müller, Michael N. Hall, Mitsugu Shimobayashi

## Abstract

Mammalian target of rapamycin complex 2 (mTORC2) is a protein kinase complex that plays an important role in energy homeostasis. Loss of adipose mTORC2 reduces lipogenic enzyme expression and *de novo* lipogenesis in adipose tissue. Adipose-specific mTORC2 knockout mice also displays triglyceride accumulation in the liver. However, the mechanism and physiological role of hepatic triglyceride accumulation upon loss of adipose mTORC2 are unknown. Here, we show that loss of adipose mTORC2 increases expression of *de novo* lipogenic enzymes in the liver, thereby causing accumulation of hepatic triglyceride and hypertriglyceridemia. Simultaneous inhibition of lipogenic enzymes in adipose tissue and liver by ablating mTORC2 in both tissues prevented accumulation of hepatic triglycerides and hypertriglyceridemia. However, loss of adipose and hepatic mTORC2 caused severe insulin resistance and glucose intolerance. Thus, our findings suggest that increased hepatic lipogenesis is a compensatory mechanism to cope with loss of lipogenesis in adipose tissue, and further suggest that mTORC2 in adipose tissue and liver plays a crucial role in maintaining whole-body energy homeostasis.

**Graphical abstract:** 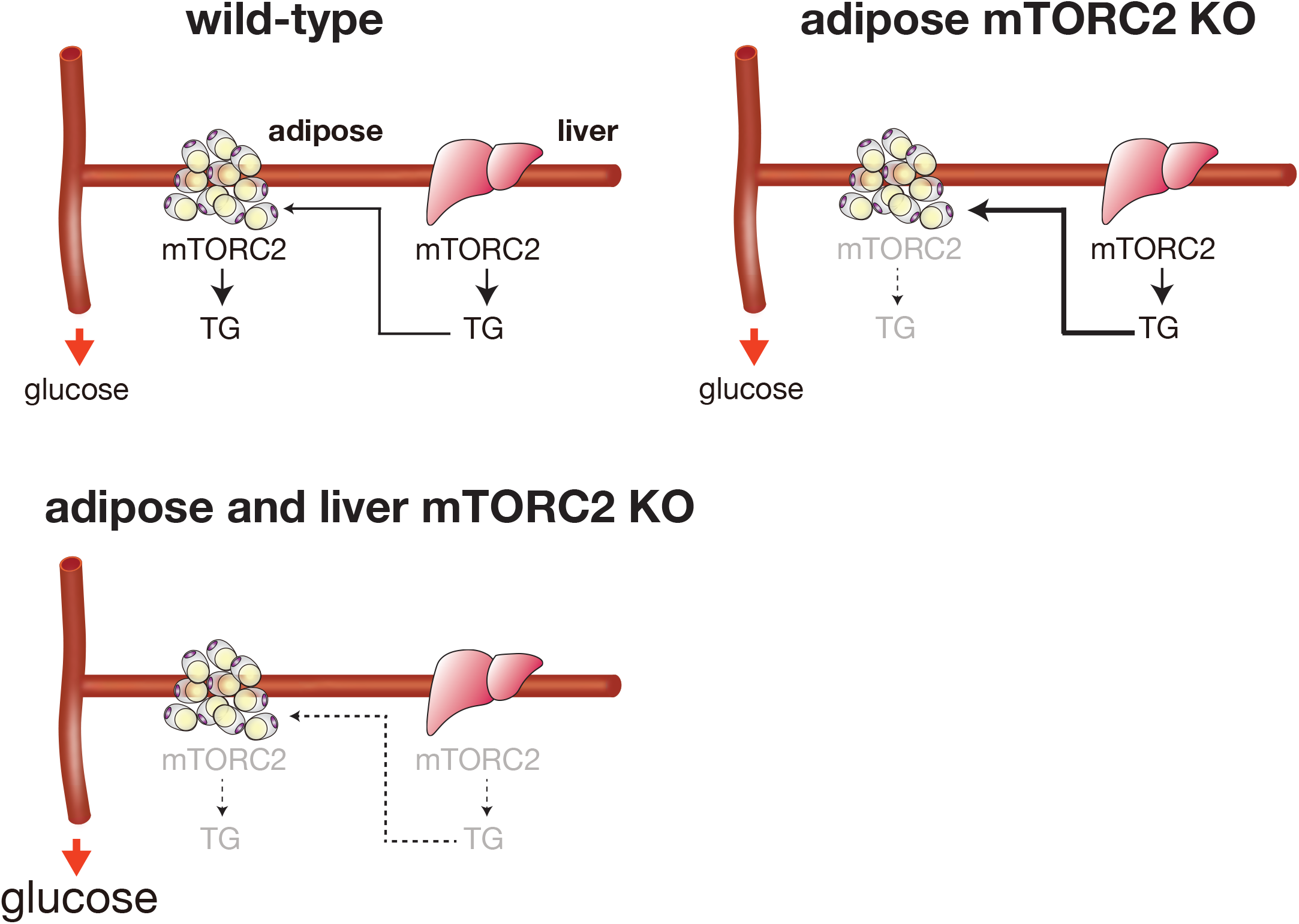

## Introduction

Organisms adapt their metabolism in response to nutrient availability. Under nutrition rich conditions, excessive energy is stored in white adipose tissue (WAT) in the form of triglyceride (TG) (1). The physiological importance of WAT is well documented in studies of patients with lipodystrophy, who fail to store a surplus of energy in WAT due to impaired WAT function (2). Although most patients with lipodystrophy appear lean, they develop disorders commonly associated with obesity, such as cardiovascular diseases, type II diabetes, non-alcoholic fatty liver disease (NAFLD), and hypertriglyceridemia.

The hormone insulin promotes energy storage. In WAT, insulin promotes the uptake of glucose, which is converted into free fatty acids (FFAs) and triglycerides (TGs) by a metabolic process defined as *de novo* lipogenesis (DNL). One of the major kinases regulating insulin-stimulated glucose uptake and DNL is mammalian target of rapamycin complex 2 (mTORC2)(3, 4). mTORC2 is a serine/threonine kinase complex of which the core components including the kinase mTOR and rapamycin-insensitive companion of mTOR (RICTOR) (5–7). The physiological role of mTORC2 in adipose tissue has been studied *in vivo* by characterizing conditional *Rictor* knockout mice (3, 8–10). It has been demonstrated that loss of adipose mTORC2 impairs the expression of genes encoding lipogenic enzymes including fatty acid synthase (FASN) and acetyl-CoA carboxylase (ACC) and thus DNL (3, 4). Mice lacking adipose mTORC2 develop hyperinsulinemia, systemic insulin resistance and hepatic TG accumulation (3, 8, 10). Surprisingly, mice lacking adipose mTORC2 mice remain glucose tolerant, suggesting presence of a compensatory mechanism that counteracts the reduced storage capacity of WAT lacking mTORC2.

Similar to WAT, loss of hepatic mTORC2 reduces the expression of the lipogenic enzymes FASN and ACC and hepatic TG synthesis, causing hypotriglyceridemia. Mice lacking hepatic mTORC2 are mildly insulin resistant and glucose intolerant (11), indicating that hepatic mTORC2 is important to maintain whole-body energy homeostasis.

In this study, we show that acute loss of mTORC2 in mature adipocytes causes mild lipodystrophy due to impaired glucose uptake and reduced expression of lipogenic enzymes. We found that the expression of hepatic lipogenic enzymes and hepatic and plasma TG levels are upregulated upon loss of adipose mTORC2. We hypothesized that the increase in hepatic DNL and TG is a mechanism to compensate for impaired WAT function and to maintain whole-body glucose homeostasis. Deletion of mTORC2 in both adipose tissue and liver blocked the expression of DNL enzymes in both organs and prevented TG accumulation in the liver and hypertriglyceridemia. However, mice lacking both adipose and hepatic mTORC2 developed severe insulin resistance and glucose intolerance. Thus, our findings suggest that increased hepatic DNL upon loss of adipose mTORC2 is a physiological response to compensate for impaired energy storge in WAT.

## Results

### Loss of mTORC2 in adult mice causes mild lipodystrophy

Previous studies have shown that mice lacking adipose mTORC2 from birth display impaired glucose uptake and reduced expression of lipogenic enzymes including FASN and ACC in WAT (3, 4). However, adipose tissue is not fully developed until six weeks of age and thus the impaired glucose uptake and DNL could be due to a developmental defect. To examine the role of mTORC2 in mature adipocytes in adult mice, we characterized inducible adipose-specific *Rictor* knockout mice (iAdRiKO: *Rictor^fl/fl^ Adipoq-CreER^T2^*) (12). *Rictor* deletion was induced by tamoxifen injections at six- to eight-week of age and loss of RICTOR and mTORC2 activity were confirmed by immunoblotting (**Figures 1A-B**). Next, we examined expression of lipogenic enzymes in WAT. Consistent with previous studies (3, 4), FASN and ACC expression were downregulated in WAT of iAdRiKO mice, compared to controls, indicating reduced DNL (**Figures 1A-B**). To examine whether loss of adipose mTORC2 in mature adipocytes also caused reduced glucose uptake, we treated iAdRiKO mice with the glucose analog 2-deoxyglucose (2DG) and insulin. Glucose uptake was reduced in WAT of iAdRiKO mice, compared to controls (**Figure 1C**). These findings confirmed previous observations that mTORC2 promotes both glucose uptake and the expression of lipogenic enzymes in mature adipocytes (3, 4). Next, we examined whether reduced glucose uptake and DNL in WAT had impact on body weight and fat mass. Similar to mice lacking adipose mTORC2 from birth (3, 8, 10), iAdRiKO mice displayed normal body weight (**Figure 1D**). However, body composition analysis revealed a ~25% reduction in fat mass and a corresponding increase in lean mass in iAdRiKO mice (**Figure 1E**). Furthermore, we found that epididymal and inguinal WAT depots, but not brown adipose tissue (BAT), were decreased in weight in iAdRiKO mice (**Figure 1F**). Taken together, these data suggest that loss of mTORC2 in mature adipocytes causes mild lipodystrophy, likely due to impaired glucose uptake and DNL in WAT.

**Figure 1.**
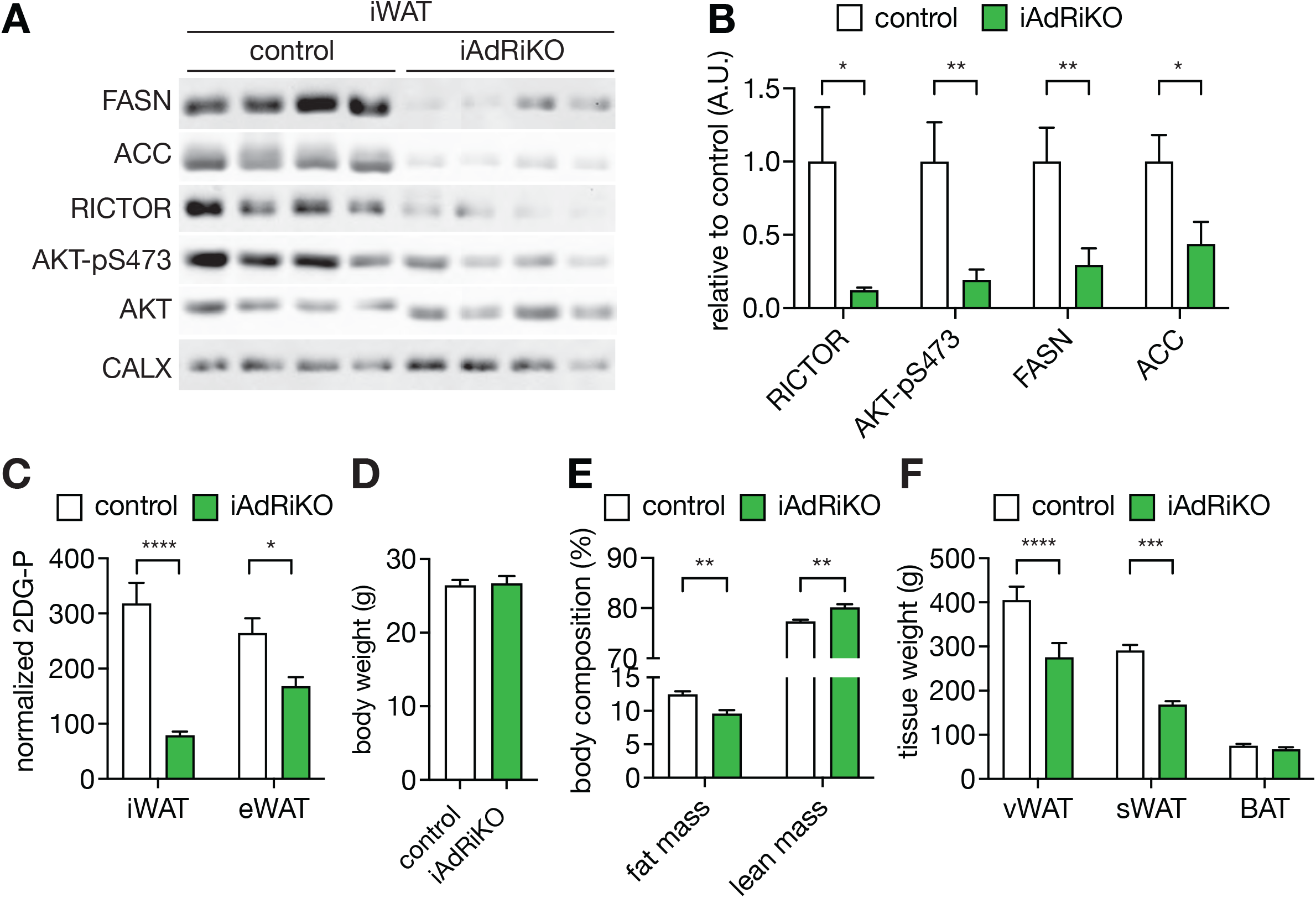
Loss of adipose mTORC2 in adult mice causes mild lipodystrophy. (A) Immunoblot analyses of inguinal WAT (iWAT) from control and iAdRiKO mice. CALX serves as a loading control. n=9 (control) and 12 (iAdRiKO). (B) Quantification of immunoblots in A. The intensities of RICTOR, FASN, ACC were normalized to CALX and the intensities of AKT-pS473 were normalized to AKT. Student’s t test, *p<0.05, **p<0.01. (C) 2-deoxyglucose (2DG) uptake in iWAT and epididymal WAT (eWAT) of control and iAdRiKO mice (n= 6). Student’s t test, **** p<0.0001, * p<0.05. (D) Body weight of control and tamoxifen inducible *Rictor* knockout (iAdRiKO) mice. n=9 (control) and 8 (iAdRiKO). (E) Body composition of control and iAdRiKO mice. n=5 (control) and 8 (iAdRiKO). Student’s t test, **p<0.01. (F) Organ weight for epididymal white adipose tissue (eWAT), iWAT, and brown adipose tissue (BAT). n=9 (control) and 8 (iAdRiKO). Student’s t test, ***p<0.001, ****p<0.0001.

### Loss of adipose mTORC2 in adult mice causes increased expression of lipogenic enzymes and TG levels in the liver

Despite reduced glucose uptake in WAT, iAdRiKO mice remained glucose tolerant (**Figure 2A**). Thus, we hypothesized that another organ takes up glucose to compensate for impaired glucose uptake in WAT. Similar to the phenotype of in mice lacking adipose mTORC2 from birth (3, 10), iAdRiKO mice displayed increased liver weight while other organs were unaffected (**Figures 2B-C**). Histological analyses revealed an increase in the number of lipid droplets in the liver of iAdRiKO mice, compared to controls (**Figure 2D**). Accumulation of hepatic TG was also confirmed by biochemical measurements, although the difference did not reach statistical significance (**Figure 2E**). Based on these observations, we hypothesized that liver compensates for impaired glucose uptake and DNL in WAT. Indeed, we observed increased expression of FASN and ACC in the liver of iAdRiKO mice (**Figures 2F-G**). Normally, the liver secretes *de novo* synthesized TGs for storage in WAT. Consistent with the increased expression of lipogenic enzymes in the liver, plasma TG levels were significantly higher in iAdRiKO mice, compared to controls (**Figure 2H**). These data suggest that loss of adipose mTORC2 increases hepatic DNL as a mechanism to compensate for impaired glucose uptake and DNL in adipose tissue.

**Figure 2.**
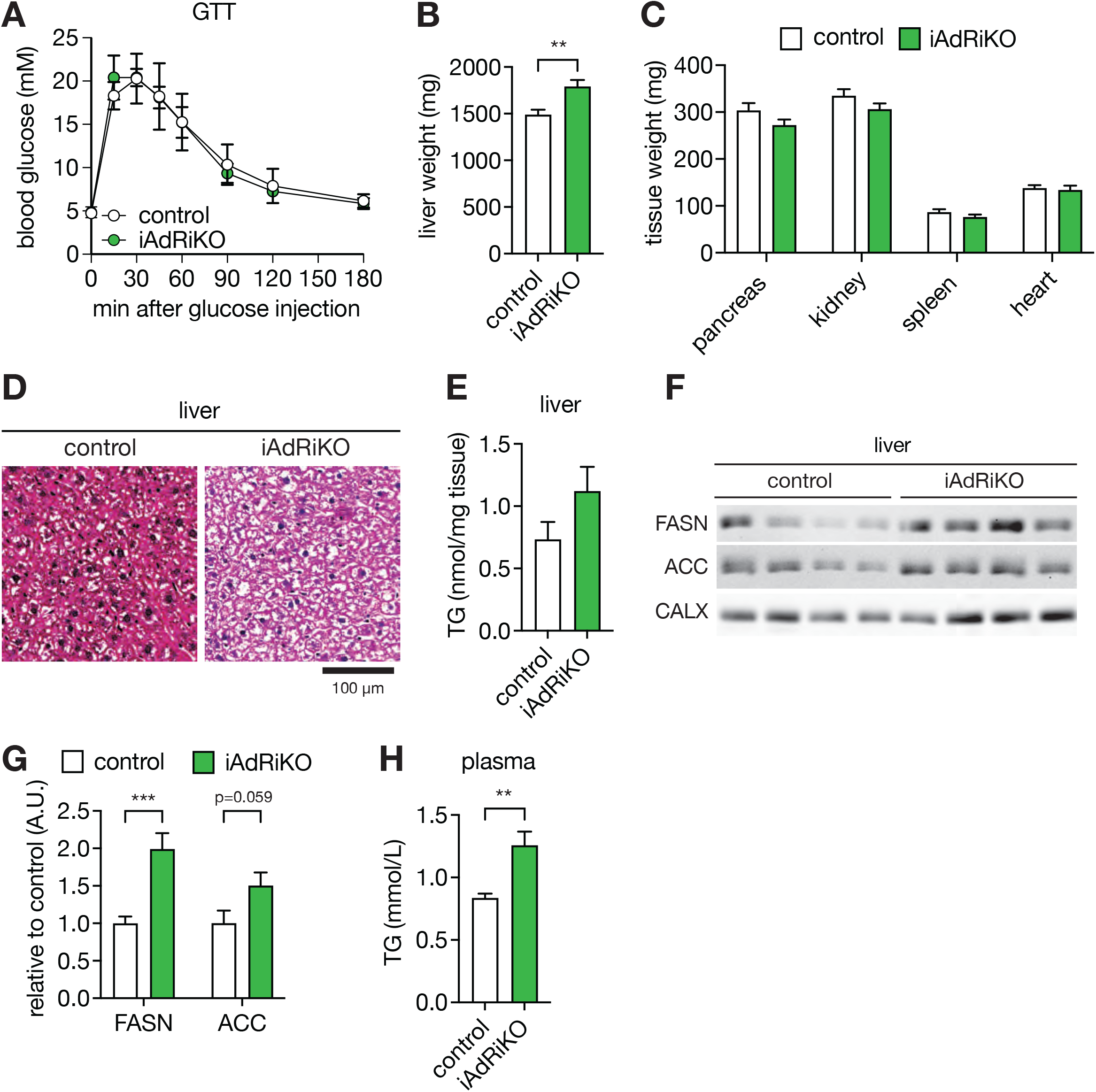
Loss of adipose mTORC2 in adult mice increases expression of lipogenic enzymes in the liver and causes hypertriglyceridemia. (A) Glucose tolerance test on control and iAdRiKO mice. n=6 (B) The liver weight of control and iAdRiKO mice. Student’s t test, **p<0.01. n=9 (control) and 8 (iAdRiKO). (C) Organ weight for pancreas, kidney, spleen and heart. n=7 (control) and 6 (iAdRiKO). (D) Hematoxylin and eosin staining of control and iAdRiKO liver. n=3 (control) and 3 (iAdRiKO). (E) Hepatic triglyceride (TG) levels in control and iAdRiKO mice. Student’s t test. n=9 (control) and 12 (iAdRiKO). (F) Immunoblot analyses of liver from control and iAdRiKO mice. CALX serves as a loading control. n=9 (control) and 12 (iAdRiKO). (G) Quantification of immunoblots in E. The intensities of FASN and ACC were normalized to CALX. Student’s t test, **p<0.01. (H) Plasma TG levels in control and iAdRiKO mice. n=15 (control) and 15 (iAdRiKO). Student’s t test, **p<0.01.

### Compensatory increase in hepatic DNL upon impaired WAT function is diminished upon loss of hepatic mTORC2

To examine the physiological role of hepatic DNL in iAdRiKO mice, we inhibited DNL in both adipose tissue and liver. mTORC2 promotes the expression of FASN and ACC also in the liver (3, 11). Thus, we generated adipose- and liver-specific double *Rictor* knockout mice (dRiKO: *Adipoq-CreER^T2^, Alb-Cre, Rictor^fl/fl^*) by crossing iAdRiKO with liver-specific *Rictor* knockout mice (LiRiKO: *Alb-Cre; Rictor^fl/fl^*). No difference in body weight was observed in dRiKO mice compared to iAdRiKO, LiRiKO, and control mice (**Figure 3A**). Similar to iAdRiKO mice, dRiKO mice displayed decreased fat mass and smaller WAT depots, compared to controls (**Figures 3B-D**). Histological analyses of WAT revealed a reduction in adipocyte diameter in WAT of iAdRiKO and dRiKO mice, indicating mild lipodystrophy (**Figures 3E-3F**). Next, we examined glucose uptake and DNL in WAT. We found that both iAdRiKO and dRiKO mice displayed reduced glucose uptake in WAT, compared to controls (**Figure 3G**). Glucose uptake was also slightly decreased in WAT of LiRiKO mice, although this difference did not reach statistical significance (**Figure 3G**). Hepatic mTORC2 ablation increases the expression of lipogenic enzymes such as FASN and ACC in WAT, although the differences did not reach statistical significance (**Figures 3H-I**). Liver weight was dereased in dRiKO mice compared to control and iAdRiKO mice (**Figure 4A**). Histological and biochemical analyses revealed a decrease in hepatic and plasma TG in dRiKO mice, compared to control and iAdRiKO mice (**Figures 4B-C**). To examine whether the decrease in hepatic and plasma TG was due to decreased expression of lipogenic enzymes in the liver, we analyzed hepatic FASN and ACC expression. Loss of hepatic mTORC2 reduced FASN and ACC expression in the liver of both LiRiKO and dRiKO mice (**Figures 4D-4E**). These data suggest that hepatic mTORC2 is required for the compensatory increase in DNL upon impaired WAT function.

**Figure 3.**
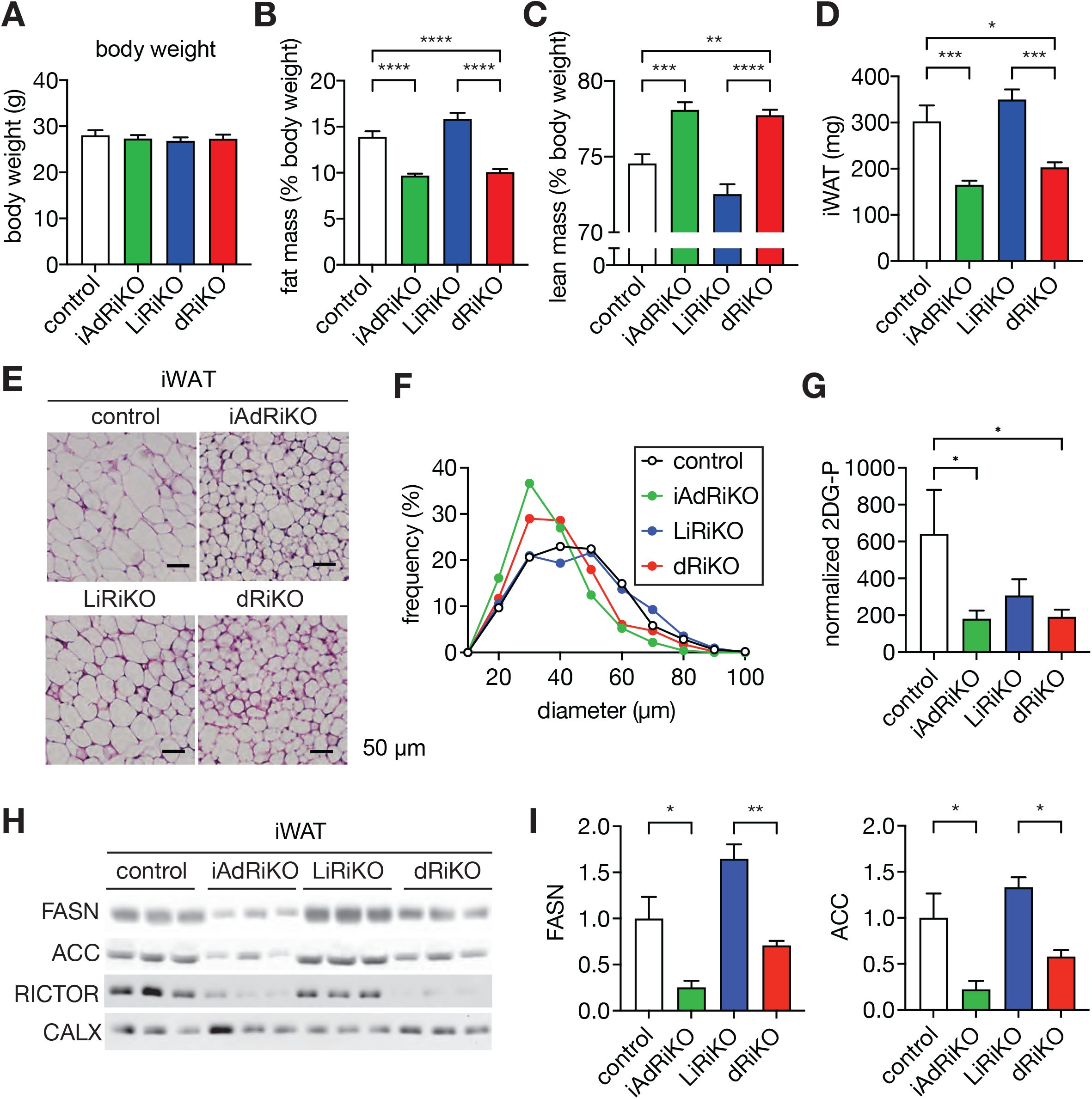
Loss of adipose and hepatic mTORC2 causes mild lipodystrophy. (A) Body weight of control, iAdRiKO, liver-specific *Rictor* knockout (LiRiKO), and adipose- and liver-specific double *Rictor* knockout (dRiKO) mice. n=9 (control), 12 (iAdRiKO), 11 (LiRiKO), and 7 (dRiKO). (B-C) Fat mass (B) and lean mass (C) of control, iAdRiKO, LiRiKO, and dRiKO mice. n=11 (control), 12 (iAdRiKO), 13 (LiRiKO), and 12 (dRiKO). One-way ANOVA, **p<0.01, ***p<0.001, ****p<0.0001. (D) iWAT weight of control, iAdRiKO, LiRiKO, and dRiKO mice. n=9 (control), 11 (iAdRiKO), 11 (LiRiKO), and 7 (dRiKO). One-way ANOVA, *p<0.05, ***p<0.001. (E) Hematoxylin and eosin staining of control, iAdRiKO, LiRiKO, and dRiKO iWAT. n=7 (control), 3 (iAdRiKO), 6 (LiRiKO), 7 (dRiKO). (F) Quantification of adipocyte diameters in the images in E. (G) 2-deoxyglucose (2DG) uptake in iWAT of control, iAdRiKO, LiRiKO and dRiKO mice. n=6 (control), 11 (iAdRiKO), 8 (LiRiKO), 12 (dRiKO). Student’s t test, * p<0.05 (H) Immunoblot analyses of iWAT from control, iAdRiKO, LiRiKO, and dRiKO mice. CALX serves as a loading control. n=9 (control), 12 (iAdRiKO), 11 (LiRiKO), 7(dRiKO). (I) Quantification of immunoblots in Figure 3H. The intensities of FASN and ACC were normalized to CALX. One-way ANOVA, *p<0.05, **p<0.01.

**Figure 4.**
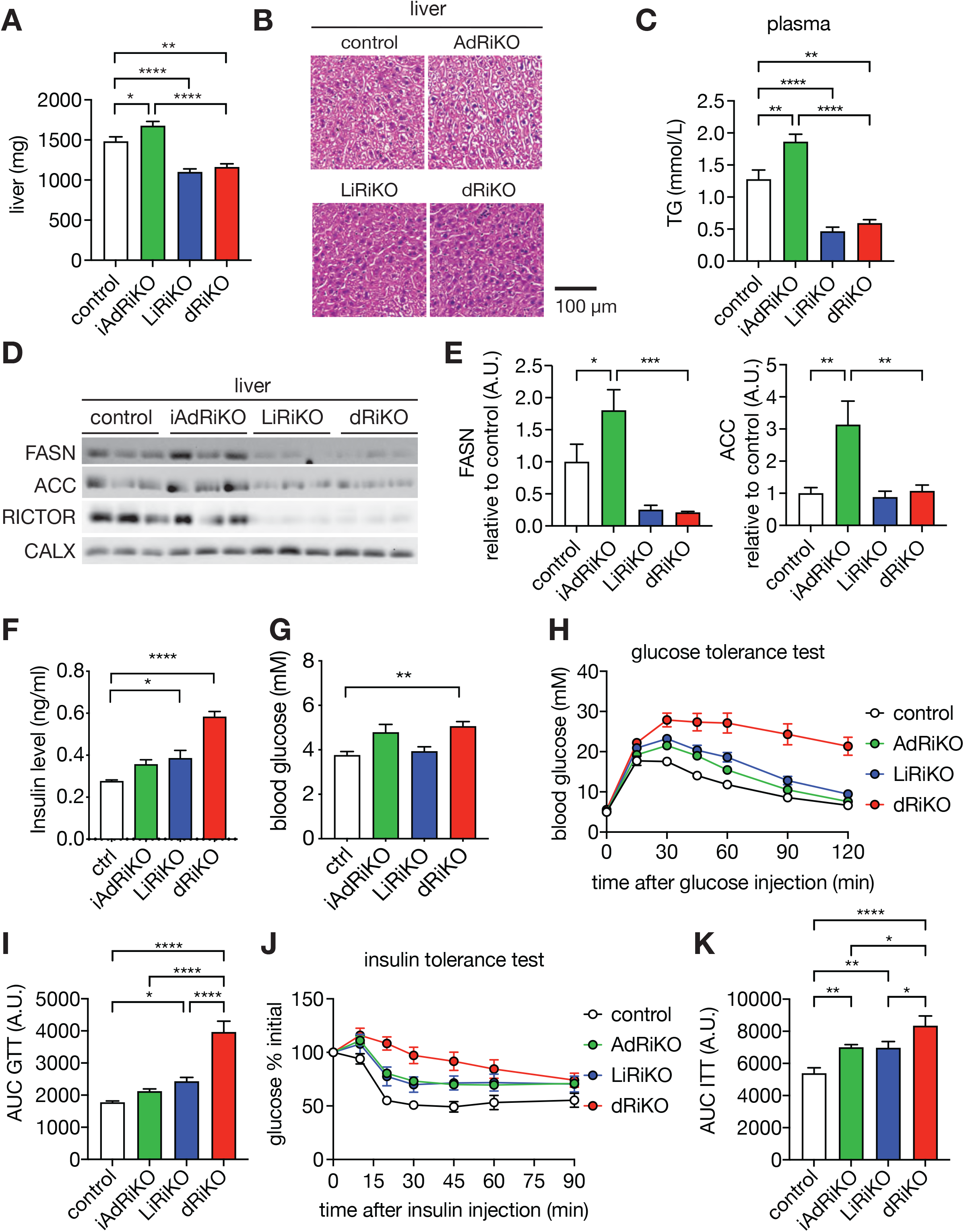
Loss of adipose and hepatic mTORC2 reduces expression of lipogenic enzymes in the liver and causes hypotriglyceridemia, but causes insulin resistance and severe glucose intolerance. (A) The liver weight of control, iAdRiKO, LiRiKO, and dRiKO mice. n=9 (control), 11 (iAdRiKO), 11 (LiRiKO), 7 (dRiKO). One-way ANOVA, *p<0.05, **p<0.01, ****p<0.001. (B) Hematoxylin and eosin staining of control, iAdRiKO, LiRiKO, and dRiKO liver. n=3 (control), 3 (iAdRiKO), 3 (LiRiKO), 3 (dRiKO). (C) Plasma TG levels in control, iAdRiKO, LiRiKO, and dRiKO mice. n=9 (control), 11 (iAdRiKO), 11 (LiRiKO), 7 (dRiKO). One-way ANOVA, **p<0.01, ****p<0.001. (D) Immunoblot analyses of liver from control, iAdRiKO, LiRiKO, and dRiKO mice. CALX serves as a loading control. n=6 (control), 6 (iAdRiKO), 6 (LiRiKO), 6 (dRiKO). (E) Quantification of immunoblots in D. The intensities of FASN and ACC were normalized to CALX. One-way ANOVA, *p<0.05, **p<0.01. (F) Plasma insulin levels of control, iAdRiKO, LiRiKO and RidKO mice after 16 hour starvation. One-way ANOVA, *p<0.05, ****p<0.0001. n= 4 (control), 5 (iAdRiKO), 5 (LiRiKO), 7 (dRiKO). (G) Blood glucose levels of control, iAdRiKO, LiRiKO and RidKO mice mice after 16 hour starvation. n= 4 (control), 5 (iAdRiKO), 5 (LiRiKO), 7 (dRiKO). One-way ANOVA, **p<0.01. (H) Glucose tolerance test on control, iAdRiKO, LiRiKO, and dRiKO mice. The mice were fasted for 14 hours and injected with glucose (2 g/kg body weight). n=7 (control), 19 (iAdRiKO), 10 (LiRiKO), 9 (dRiKO). (I) Area under the curve (AUC) of the blood glucose curve in H. One-way ANOVA. *p<0.05, ****p<0.0001. (J) Insulin tolerance test on control, iAdRiKO, LiRiKO, and dRiKO mice. The mice were fasted for 6 hours and injected with insulin (0.75 U/kg body weight). n=18 (control), 19 (iAdRiKO), 11 (LiRiKO), 9 (dRiKO). (K) Area under the curve (AUC) of the blood glucose curve in E. One-way ANOVA. *p<0.05, **p<0.01, ****p<0.0001.

### Simultaneous loss of adipose and hepatic mTORC2 causes severe insulin resistance and glucose intolerance

Next, we investigated whole body energy homeostasis upon loss of both adipose and hepatic mTORC2 (dRiKO). Fasting plasma insulin and glucose level were both significantly increased in dRiKO mice compared to control mice (**Figures 4F-G**). Since glucose is the major source for DNL in both adipose tissue and liver, we assessed whole body glucose homeostasis by performing a glucose tolerance test. As previously reported, LiRiKO mice were slightly glucose intolerant (11) while iAdRiKO and control mice were glucose tolerant (**Figures 4H-I**). In contrast, dRiKO mice displayed severe glucose intolerance, the severity of which is synergistic compared to the effect observed in iAdRiKO and LiRiKO mice (**Figures 4H-I**). Next, we performed an insulin tolerance test to determine systemic insulin sensitivity. Consistent with previous studies (3, 8, 9, 11), iAdRiKO and LiRiKO mice were insulin resistant compared to control mice (**Figures 4J-K**). dRiKO mice displayed severe insulin resistance. Taken together, our findings suggest that adipose and liver mTORC2 are indispensable to maintain whole body glucose homeostasis and that simultaneous loss of both hepatic and adipose mTORC2 signaling synergistically causes severe diabetes.

## Discussion

In this study, we highlight the physiological importance of adipose and hepatic mTORC2 in mediating whole body lipid and glucose homeostasis (**graphical abstract**). Loss of mTORC2 in mature adipocytes impaired adipose DNL, caused mild lipodystrophy, and increased hepatic DNL and TG synthesis (**graphical abstract**). Loss of hepatic mTORC2 in mice lacking adipose mTORC2 blocked hepatic TG accumulation and DNL and caused severe diabetes (**graphical abstract**). Thus, our findings suggest that increased hepatic TG accumulation and DNL in mice with impaired WAT function (e.g. mTORC2 loss) is a physiological response to compensate for impaired WAT function.

AKT is the best characterized mTORC2 substrate. It was previously reported that loss of adipose AKT1 and AKT2 causes lipodystrophy and hepatic TG accumulation (13). Adipose-specific AKT1 and AKT2 knockout (Adipo-AKT KO) mice are insulin resistant but glucose tolerant. It is unknown why Adipo-AKT KO mice remain glucose tolerant. Similar to Adipo-AKT KO mice, we show that mice lacking adipose mTORC2 displayed lipodystrophy, hepatic TG accumulation and systemic insulin resistance but remained glucose tolerant. Accumulation of hepatic TG in mice lacking adipose mTORC2 is due to increased expression of lipogenic enzymes in the liver. Blocking hepatic DNL and TG accumulation by ablation of hepatic mTORC2 causes severe insulin resistance and glucose intolerance, suggesting that increased hepatic DNL and TG accumulation in iAdRiKO mice is a compensatory mechanism ensuring whole-body glucose homeostasis. Considering that AKT 1 and AKT2 are mTORC2 substrates, our findings suggest that TG accumulation observed in Adipo-AKT KO mice may also be a compensatory mechanism mediating whole-body glucose homeostasis.

Mice lacking adipose mTORC2 from birth displayed no change in fat mass (3, 8), but the current study showed a reduction in fat mass upon loss of adipose mTORC2 in adult mice. How can this difference be explained? It has been demonstrated that adiponectin is expressed in adipocyte precursors in WAT (14) and that *Adipoq* expression is already detectable in WAT at embryonic day 17.5 (15). Although the deletion of RICTOR in adipocyte precursors needs to be confirmed, loss of mTORC2 during early adipogenesis may allow mice to adapt to maintain adipose function.

Hepatic TG accumulation is a common symptom observed in patients and mouse models of lipodystrophy (2). Since hepatocytes are not specialized to store high quantities of TG, patients and mice with impaired WAT develop NAFLD, a chronic liver disease that can progress to cirrhosis and hepatocellular carcinoma. To treat NAFLD, DNL inhibitors are currently being tested in clinical trials (16). However, based on our finding that dRiKO mice displayed reduced hepatic DNL but severe diabetes, targeting hepatic DNL may not be recommended to treat NAFLD in patients with impaired WAT function.

The immunosuppressive drug rapamycin has been shown to extend lifespan and prevent age-related diseases, including neurogenerative disease and cancer, by inhibiting the mTORC1. However, it has been shown that rapamycin treatment causes insulin resistance and glucose intolerance (17, 18). A previous study demonstrated that these rapamycin-induced adverse effects are due mainly to inhibition of hepatic mTORC2 (19). However, in this previous study, the contribution of mTORC2 loss in both adipose tissue and the liver was not investigated. Our observations in dRiKO mice suggest that inhibition of both adipose and hepatic mTORC2 contribute to the pathogenesis of rapamycin-induced insulin resistance and glucose intolerance. Thus, our findings stress the necessity of developing selective mTORC1 inhibitors to extend lifespan and to target age-related disease (20, 21).

## Materials and Methods

### Mice

Tamoxifen-inducible adipose-specific *Rictor* knockout (iAdRiKO:*Rictor^fl/fl^, adipoq-CreER^T2^*) and liver specific-Rictor knockout (LiRiKO: *Rictor^fl/fl^, alb-Cre*) mice were described previously (11, 22). iAdRiKO and LiRiKO mice were crossed to generate adipose- and liver-specific double *Rictor* knockout (dRiKO: *Rictor^fl/fl^, adipoq-CreER^T2^, alb-Cre*) mice. For iAdRiKO and dRiKO mice, *Rictor* knockout was induced by *i.p. injection of* 1 mg/mouse tamoxifen (Sigma-Aldrich) in corn oil (Sigma-Aldrich) for 5 days. Littermate *Cre* negative animals were used as a control. Control mice were also treated with tamoxifen. Mice were housed at 22 °C in a conventional facility with a 12 hour light/dark cycle with unlimited access to water and normal chow diet (KLIBA). Only male mice between 6 and 12 weeks of age were used for experiments. Body composition was measured by nuclear magnetic resonance imaging (Echo Medical Systems).

### Immunoblots

Isolated tissues were snap frozen in liquid nitrogen and stored at −80 °C. Frozen tissues were homogenized in a lysis buffer containing 100 mM Tris (Merck) pH7.5, 2 mM EDTA (Sigma-Aldrich), 2 mM EGTA (Sigma-Aldrich), 150 mM NaCl (Sigma-Aldrich), 1% Triton X-100 (Fluka), cOmplete inhibitor cocktail (Roche) and PhosSTOP (Roche). Protein concentration was determined by Bradford assay (Biorad), and equal amounts of protein were separated by SDS-PAGE, and transferred onto nitrocellulose membranes (GE Healthcare). Antibodies used in this study were as follows: RICTOR (Cat#2140) AKT (Cat#4685 or Cat#2920), AKT-pS473 (Cat#4060), FASN (Cat#3189), and ACC (Cat#3662) from Cell Signaling Technology, and CALNEXIN (Cat#ADI-SPA-860-F) from Enzo Life Sciences.

### Body composition measurement

Body composition was measured by nuclear magnetic resonance imaging (Echo Medical Systems).

### 2-Deoxyglucose uptake assay

Mice were fasted for five hours, then injected i.p. with Humalog insulin (Lilly; 0.75 U/kg body weight), followed 10 minutes later with an injection of 2-deoxyglucose (Sigma-Aldrich; 32.8 μg/g body weight). Tissues were collected 20 minutes after administration of 2-deoxyglucose. Tissues were lysed in 10 mM Tris-HCl, pH 8.0, by boiling for 15 minutes. 2-Deoxyglucose-6-phosphate (2DGP) was measured using a Glucose Uptake-Glo Assay Kit (Promega) following the manufacturer’s instructions.

### Histology

Tissues were fixed in 4% formalin (Biosystems), embedded in paraffin (Biosystems), and sliced into 4 μm thick sections. Tissue sections were stained with Hematoxylin (Sigma-Aldrich) and eosin (Waldeck), and imaged by a conventional microscope (Zeiss). Adipocyte diameters were quantified with FIJI by using the Adiposoft plugin (23).

### Triglyceride measurement

50-100 mg liver tissues were homogenized by a dounce homogenizer in 5% NP-40 in ddH_2_O. The homogenates were slowly heated up to 90 °C for 5 minutes and cooled down to room temperature. This heating-cooling cycle was repeated once more. The homogenates were centrifuged at 14’000 g for 2 min and the supernatant was used to measure triglyceride using a triglyceride quantification kit (Abcam) by following the manufacture’s instruction. The hepatic TG levels were normalized to liver weights. Plasma TG levels were measured by a biochemical analyzer (Cobas c III analyser, Roche) or a triglyceride quantification kit (Abcam).

### Fasting insulin measurement

Mice were fasted for 16 hours and blood samples were collected to determine fasting insulin and blood glucose levels. Plasma insulin levels were measured by ultrasensitive mouse insulin ELISA kit (Crystal Chem) according to the manufacturer’s instructions.

### Insulin and glucose tolerant test

For the insulin and glucose tolerance tests, mice were fasted for 6 hours or overnight, respectively. Insulin Humalog (Lilly, i.p. 0.5 U/kg body weight) for insulin tolerance test or glucose (Sigma-Aldrich, 2 g/kg body weight) for glucose tolerance test was i.p. administered. Blood glucose was measured with a blood glucose meter (Accu-Check, Roche). Plasma insulin levels were measured by ultrasensitive mouse insulin ELISA kit (Crystal Chem) according to the manufacturer’s instructions.

### Study Approval

All animal experiments were performed in accordance with federal guidelines for animal experimentation and were approved by the Kantonales Veterinäramt of the Kanton Basel-Stadt.

### Statistics

Sample size was chosen according to our previous studies and published reports in which similar experimental procedures were described. All data are shown as the mean ± SEM. Sample numbers are indicated in each figure legend. To determine the statistical significance between 2 groups, an unpaired two-tailed Student’s t-test was performed. For more than 3 groups, one-way ANOVA was performed. For insulin tolerance test and glucose tolerance test, two-way ANOVA was performed. All statistical analyses were performed using GraphPad Prism 9 (GraphPad Software). A *p* value of less than 0.05 was considered statistically significant.

## Acknowledgements

We thank Stefan Offermanns (MPI-HLR, Germany) for mouse, Cécile Aude Pfaff (Biozentrum) for technical supports, Geert Carmeliet (KU Leuven), and Karen Moermans (KU Leuven) for supports for histology. We acknowledge support from the Swiss National Science Foundation (project 179569 and NCCR 182880 to MNH and 161510 to MS), European Foundation of the study of Diabetes/Novo Nordisk Foundation (MS), KU Leuven internal fund (MS), The Louis Jeantet Foundation (MNH), and the Canton of Basel (MNH).

## Contributions

ICF, MNH and MS conceived the project and designed the experiments. ICF, DW, CM, and MS performed experiments and analyzed data. ICF, MNH and MS wrote the manuscript.

